# Exercise Attenuates Sickness Behavior And Protects Against Dopaminergic Impairment Induced By Neuroinflammation

**DOI:** 10.1101/2021.02.05.429925

**Authors:** Ana Cristina de Bem Alves, Ananda Christina Staats Pires, Ana Elisa Speck, Samantha Cristiane Lopes, Débora da Luz Scheffer, Hiago Murilo Melo, Rui Daniel Prediger, Roberta de Paula Martins, Alexandra Latini, Aderbal S Aguiar

## Abstract

Neuroinflammation affects dopamine metabolism and produces a set of symptoms known as sickness behavior, including fever, anhedonia, anorexia, weight loss, decreased sociability and mobility, and cognitive impairment. Motor and cognitive impairments related to sickness behavior are associated with dopamine (DA) metabolism imbalance in the prefrontal cortex. Lipopolysaccharide (LPS) administration induces neuroinflammation and causes sickness behavior in mice, while physical exercise has anti-inflammatory properties and may attenuate sickness behavior and DA impairment. We investigated the effect of exercise on DA levels and sickness behavior induced by LPS in mice. Adult Swiss male mice (8–10 weeks, 47.1 ± 0.7 g, n=495) performed six weeks of voluntary exercise in free-running wheels (RW group) or had the blocked wheel in their cages (sedentary, SED group). After six weeks of exercise, both groups received an intraperitoneal injection (i.p.) of either saline (SAL) or LPS (0.33 mg/kg, i.p.). All animals were submitted to behavioral tests for sickness behavior assessment (fatigue, locomotion, anhedonia, and social interaction). Neuroinflammation markers and DA metabolism were assessed in the prefrontal cortex. LPS administration provoked anorexia, body weight loss, impaired motor function, social withdrawal, and anhedonia. This sickness behavior was accompanied by reduced cortical DA metabolism and its metabolite, 3,4-dihydroxyphenylacetic acid (DOPAC). Neuroinflammation was confirmed through increased levels of the proinflammatory cytokines IL-1β and IL-6. Inflammation was also confirmed in the blood by an increased content of IL-1β. Physical exercise intervention prevented animals from neurochemical, biochemical, and behavioral alterations. These findings provide new evidence of physical exercise’s potential as an environmental approach to treating neuroinflammatory conditions.

## INTRODUCTION

Normal levels of inflammatory cytokines are essential for brain neurocircuitry maintenance [1]. However, exacerbated and chronic inflammation and excess of inflammatory cytokines affect neuronal integrity and the monoamine neurotransmitter system in the brain, producing a range of behavioral disturbances consistent with dopamine (DA) function changes [2]. Sickness behavior is a set of symptoms induced by increased inflammatory cytokines due to viral infections, neuroinflammation, or even immune trauma [3–8]. Sickness behavior’s primary symptomatology is fever, anhedonia, anorexia, weight loss, decreased sociability, mobility, and cognitive impairment [9, 10]. Although sickness behavior has been described as a conservation-withdrawal behavioral state due to metabolic constraints [11], its symptoms impair general activity, quality of life, daily and professional activities [10]. Motor and cognitive disturbances of sickness behavior are associated with DA metabolism imbalance in the prefrontal cortex [12–16].

Lipopolysaccharide (LPS) is an endotoxin present in the outer membrane of Gram-negative bacteria [17] that increases the concentration of proinflammatory cytokines interleukin (IL)-1β, IL-6, and tumor necrosis factor-α (TNF-α) in the central nervous system (CNS) [8, 10, 18]. Neuroinflammation is a common mechanism in the pathogenesis of several neurodegenerative and psychiatric diseases, *e.g.,* Alzheimer’s disease and Parkinson’s disease [16, 19, 20]. Proinflammatory cytokines may affect multiple aspects of DA neurotransmission, including DA synthesis, reuptake, or DA packing and release, reducing DA function [1]. Basal ganglia-mediated fatigue and decreased mobility are resistant to treatment with classical stimulant medications that increase dopamine transporter (DAT) mediated DA release and/or block DA reuptake, indicating cytokine effects on DA function may impair or avoid the mechanism of action of pharmacological treatment [21].

Regular exercise contributes to brain health maintenance [22–26] through reduction of the inflammatory response [27, 28], improvement of dopaminergic neurotransmission [29], neurogenesis, cell survivor, angiogenesis [30], mitochondrial morphology [31], and blood-brain barrier integrity [31]. However, exercise effects on LPS induced-neuroinflammation and sickness behavior are contradictory [32–35]. The literature points out conflicting data on TNF-α response in exercised LPS-treated animals [36, 37]. Because these studies were carried out on a treadmill, we evaluated the effect of a light-intensity voluntary exercise protocol using running wheels (RW). Exploring the effects of physical exercise on sickness behavior will provide insights about complementary strategies to pharmacological interventions in the treatment of neuroinflammatory diseases [10, 38]. Further understanding of how neuroinflammation affects DA metabolism will enhance our knowledge of neuroinflammatory conditions’ behavioral consequences.

## MATERIALS AND METHODS

### Animals and running wheel (RW)

Adult Swiss male mice (8–10 weeks, 47.1 ± 0.7 g, n=495) from the Central Facility of Federal University of Santa Catarina (UFSC) were used. The UFSC Animal Care and Use Committee (IACUC) approved the research (protocols IACUC-PP10616 and PP10519). The animals remained in the mouse vivarium of the Biochemistry Department of UFSC. They were housed in polypropylene collective cages (38 · 32 · 17 cm) lined with wood shavings, inside ventilated cabinets (Alesco Ind, Monte Mor, SP, Brazil) under a controlled environment (12h light-dark cycle, lights on at 7:00 h, room temperature 21±1°C). The access to commercial food (3.9 kcal/g, Nuvilab CR1, Nuvital Nutrientes S/A, Colombo, PR, Brazil) and tap water was *ad libitum*.

First, the animals were divided into sedentary (SED) and exercise groups (RW). Running mice were housed in individual polypropylene cages (27 × 18 × 13 cm) equipped with RW (4½”, Super Pet, USA) to stimulate voluntary running [28, 39, 40]. Control SED animals were also isolated and had access to a “locked” RW (not spinning). Mice that performed 2 km/day in the RW for two weeks were selected to the exercise group (RW) and performed four more weeks of exercise. Running distance was measured every 24 hours at the same time (19:00 h). The free and locked wheels remained in the same location in the cage [28, 39, 40]. Wheels blockade reduces the environmental enrichment bias of running wheels, and, therefore, the effects observed derive from physical exercise only. Animals that did not meet RW or SED group criteria were allocated to other experiments (about 70%). The allocation for experimental groups was random. For each test, the experimental unit was an individual animal.

### LPS administration and sickness behavior

After six weeks of exercise, RW and SED groups received an intraperitoneal injection (volume 10 mL/kg) of either LPS 0.33 mg/kg (serotype 0127: B8, Sigma) to induce neuroinflammation [7, 41–43] or vehicle (NaCl 0,9%). We monitored running performance for another week to indicate fatigue, body mass, and food intake. Treatments were administered at 9:00-10:00h to conduct behavioral tests (open-field, splash test, and social interaction) after 4 hours (13:00-14:00h). No animals died during the experimental protocol until euthanasia. Animals were immediately euthanized (cervical dislocation) after behavioral tests. Blood and prefrontal cortex were collected for ELISA (enzyme-linked immunosorbent assay) and HPLC (high-performance liquid chromatography) analysis. Blood samples were centrifuged at 5000 rpm for 5 min for serum collection. The serum and prefrontal cortex were stored at −80°C until biochemical analysis.

The locomotor activity of animals was assessed on the open-field test. Each animal could freely explore a circular apparatus (60 cm diameter) for 5 minutes in a 10 lux lighting test room [44]. The total distance (m), rearings (number and duration), and speed (maximal and average speed, m/min) were measured using the ANY-maze™ software. The splash test evaluates the self-care and anhedonic-like behaviors of animals. Sucrose 10% was sprinkled in the back of animals [45], and grooming (frequency and duration) was manually measured for 5 minutes. The social interaction test analyzes socialization. A female mouse (male matched-age) was placed in the center of the open field with the SAL- or LPS-treated male. The interaction was manually timed for 5 minutes [46].

### Enzyme-linked immunosorbent *assay* (*ELISA*)

The prefrontal cortex was homogenized into five volumes of tris 10 mM solution containing 1 mM ethylenediamine tetra-acetic acid, triton 1%, protease inhibitors aprotinin 1μ/mL, chemostatin 1 μ/mL, leupeptin 1 μ/mL, and 1 μ/mL of phenylmethylsulfonyl fluoride. After, the sample was centrifuged (14.000 RPM, 10 minutes, 4°C), and the supernatant was used for the quantification of IL-1β e IL-6 cytokines using commercial kits (R&D System^®^).

### High-performance liquid chromatography (HPLC)

The prefrontal cortex was sonicated (5 minutes) in 10 volumes of 0.1 N perchloric acid/0.02% sodium metabisulfite. DA and its metabolite 3,4-dihydroxyphenylacetic acid (DOPAC) levels were determined by analyzing 20 μL of the supernatant in a system composed of a mobile phase containing sodium phosphate (90 mM), citric acid (50 mM), sodium heptane sulfonate (1.7 mM), EDTA (48 μM), 10% acetonitrile and pH 3.0 with a flow of 0.3 mL/minute in a column 120×2.0 mm C18 (Synergi Hydro). Under these conditions, monoamines’ retention time and their metabolites were approximately 2.7 minutes for DA and 4.2 minutes for DOPAC. The concentrations of DA and DOPAC were determined by an electrochemical detector (Waters 2465) on the HPLC (Alliance e2695, Waters, Milford, USA) and calculated as ng/mg protein [47].

### Statistics

Ten independent experiments were carried out. Data are presented as mean±SEM in graphs built using the GraphPad Prism version 5.00 for Windows, GraphPad Software, San Diego California USA (www.graphpad.com).

Statistical analyzes were performed according to an intention-to-treat principle using StatSoft, Inc. (2007). STATISTICA (data analysis software system), version 8.0. www.statsoft.com. Unicaudal two-way analysis of variance ANOVA was used to evaluate open field, splash test, social interaction, IL-6, IL-1β, DA and DA/DOPAC ratio, followed by Newman-Keuls post hoc test. ANOVA with repeated measures evaluated the evolution of running distance, body weight, and food intake, followed by the Bonferroni post hoc test. The differences were considered significant when *p* < 0.05.

Effect sizes (Cohen’s partial eta-square) were calculated for between-group changes, where a Cohen’s was used for ANOVA, defined as 0.01 small, 0.09 medium, and 0.25 large.

### Data availability

Datasets generated and analyzed during the current study are available from the corresponding author on reasonable request.

## RESULTS

### Physical exercise attenuates LPS-induced sickness behavior in mice

Swiss mice had a 22.6 % ± 5.6 adherence to the RW, and the runners performed 2.4 ± 0.9 km/day. There was a significant increase in the running distance from day 1, reaching a plateau on day 10 [*F* (41,8) = 3.8, *p* < 0.05, Figure 1A]. LPS decreased running distance within six days after LPS treatment [*F* (5,1) = 4.4, *p* < 0.05, Figure 1A].

**Figure 1.**
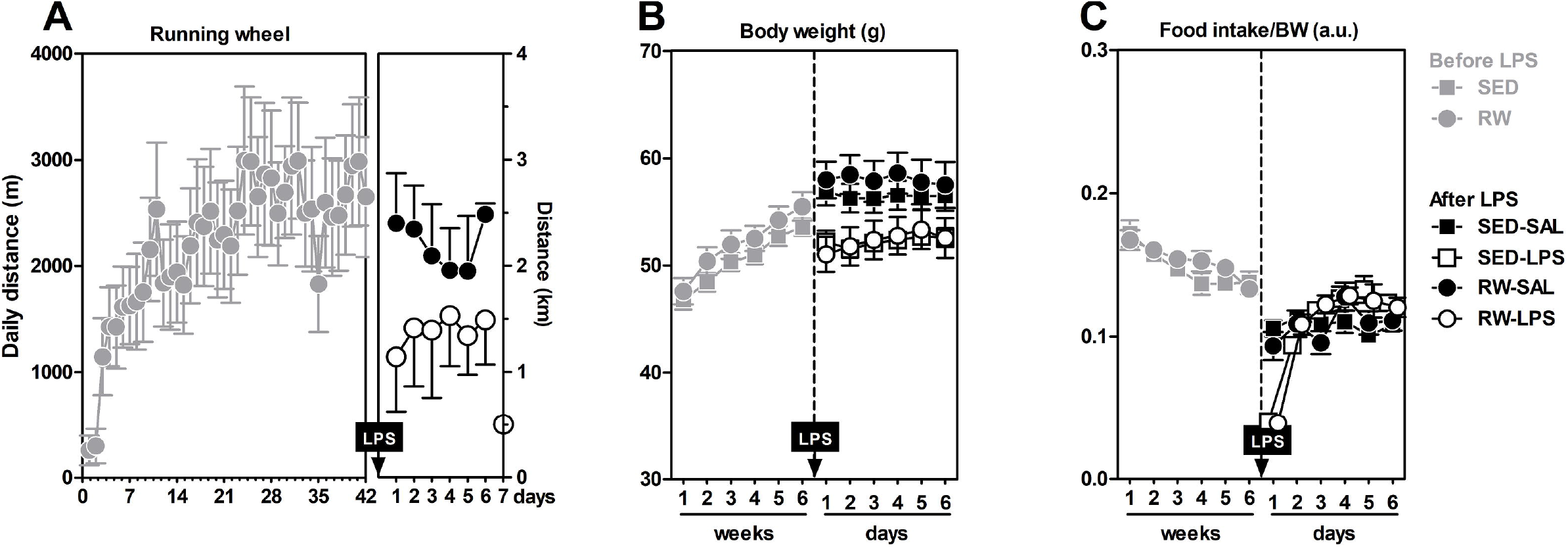
LPS (0.33 mg/kg, i.p.) induced fatigue in the running wheels (A) and loss of weight (B), and food intake (C) within one week after intoxication. Data expressed as mean ± SEM for 9-10 independent experiments. LPS – Lipopolysaccharide. RW – running wheel. SAL – saline. SED – sedentary.

There was a significant increase in the body mass of animals throughout the six weeks before treatment [*F* (5,245) =118.7, *p* < 0.05, Figure 1B], with no difference between exercised and sedentary animals. However, LPS treatment decreased the body weight [*F* (1,47) = 11.7, *p* < 0.05, Figure 1B] and food intake [*F* (1,43) = 7.3, *p* < 0.05, Figure 1C] in both SED and RW mice. However, the decrease in food intake returned to control levels (SAL) on the third day after LPS treatment, with no difference between SED and RW.

LPS decreased locomotor activity [*F* (1,28) = 11.08, *p* < 0.05, η^2^ = 0.30, Figure 2A] as well as number [*F* (1,28) = 26.47, *p* < 0.05, Figure 2B] and time [*F* (1,28) = 23.48, *p* < 0.05, Figure 2C] of rearings in the open field with large Cohen’s effect sizes (η^2^ = 0.48). The exercise did not protect against LPS-induced motor impairments (Figure 1A), but partially recovered the rearings (Figure 2B-C). LPS also decreased number [*F* (1,28) = 15.13, *p* < 0.05, Figure 2D] and time [*F* (1,28) = 7.73, *p* < 0.05, Figure 2E] of grooming in the splash test with moderate Cohen’s effect size (η^2^ = 0.23). Exercise was effective in recovering self-care of animals (Figure 2D-E). In addition, the large (η^2^ = 0.52) social isolation in SED animals treated with LPS was partially reversed by exercise [*F* (1,28) = 27.27, *p* < 0.05, Figure 2F].

**Figure 2.**
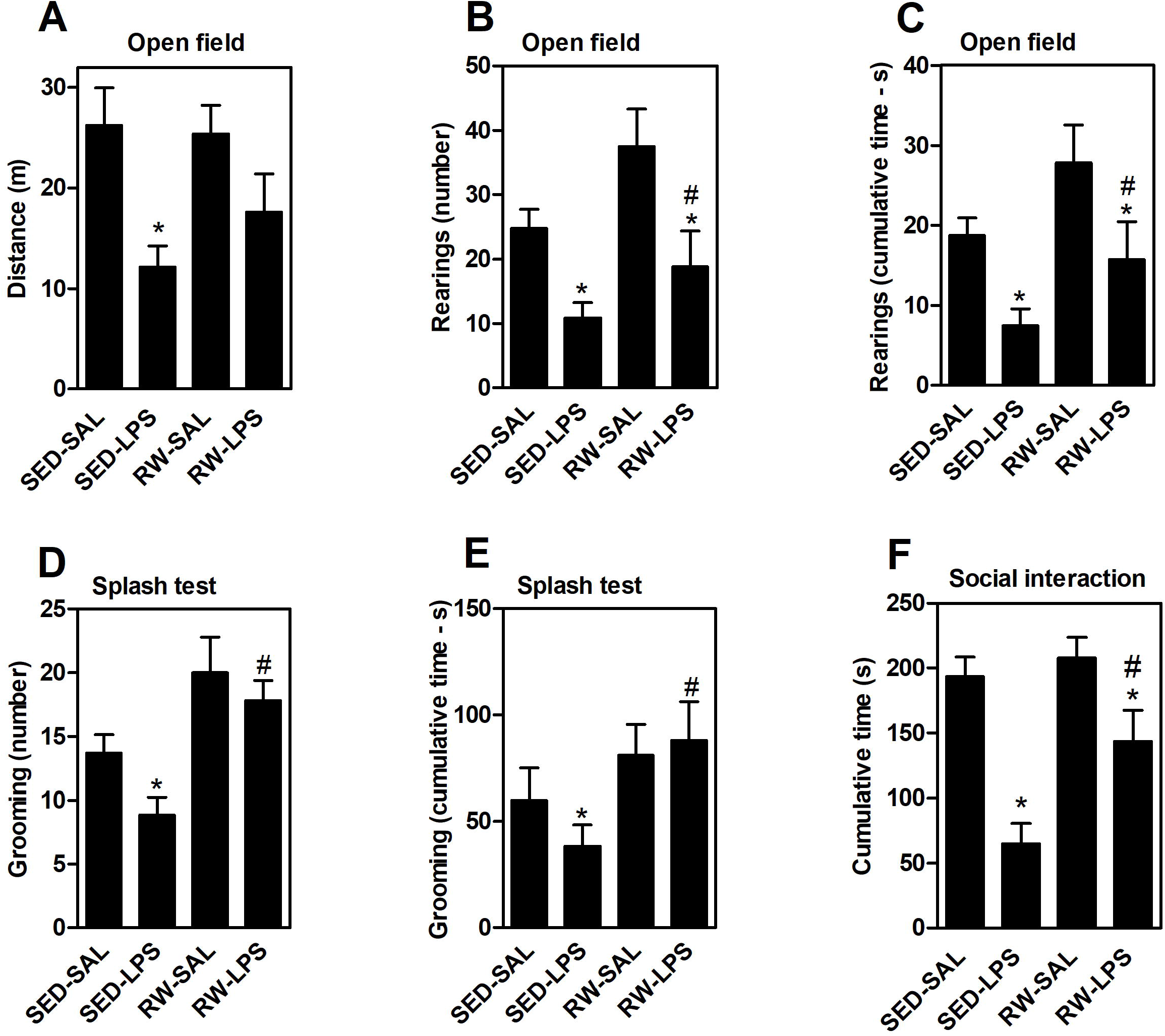
LPS (0.33 mg/kg, i.p.) induced sickness behavior in mice: motor impairment (A-C), anhedonia (D-E), and social isolation (F). Long-term running partially reversed these losses. Data expressed as mean ± SEM for 9-10 independent experiments. * P<0.05 from SAL (ANOVA two-way). # P<0.05 from SED (ANOVA two-way). LPS – Lipopolysaccharide. RW – running wheel. SAL – saline. SED – sedentary.

### Physical exercise attenuates LPS-induced cortical neuroinflammation and DA depletion

LPS increased serum [*F* (1,19) = 7.86, *p* < 0.05, η^2^ = 0.67, Figure 3A] and cortical levels of IL-1β [*F* (1,19) = 3.93, *p* < 0.05, η^2^ = 0.19, Figure 3B], with no effects of exercise. LPS also increased cortical IL-6 levels in the prefrontal cortex, which was attenuated by exercise [*F* (1,19) = 4.80, *p* < 0.05, η^2^ = 0.23, Figure 3C].

**Figure 3.**
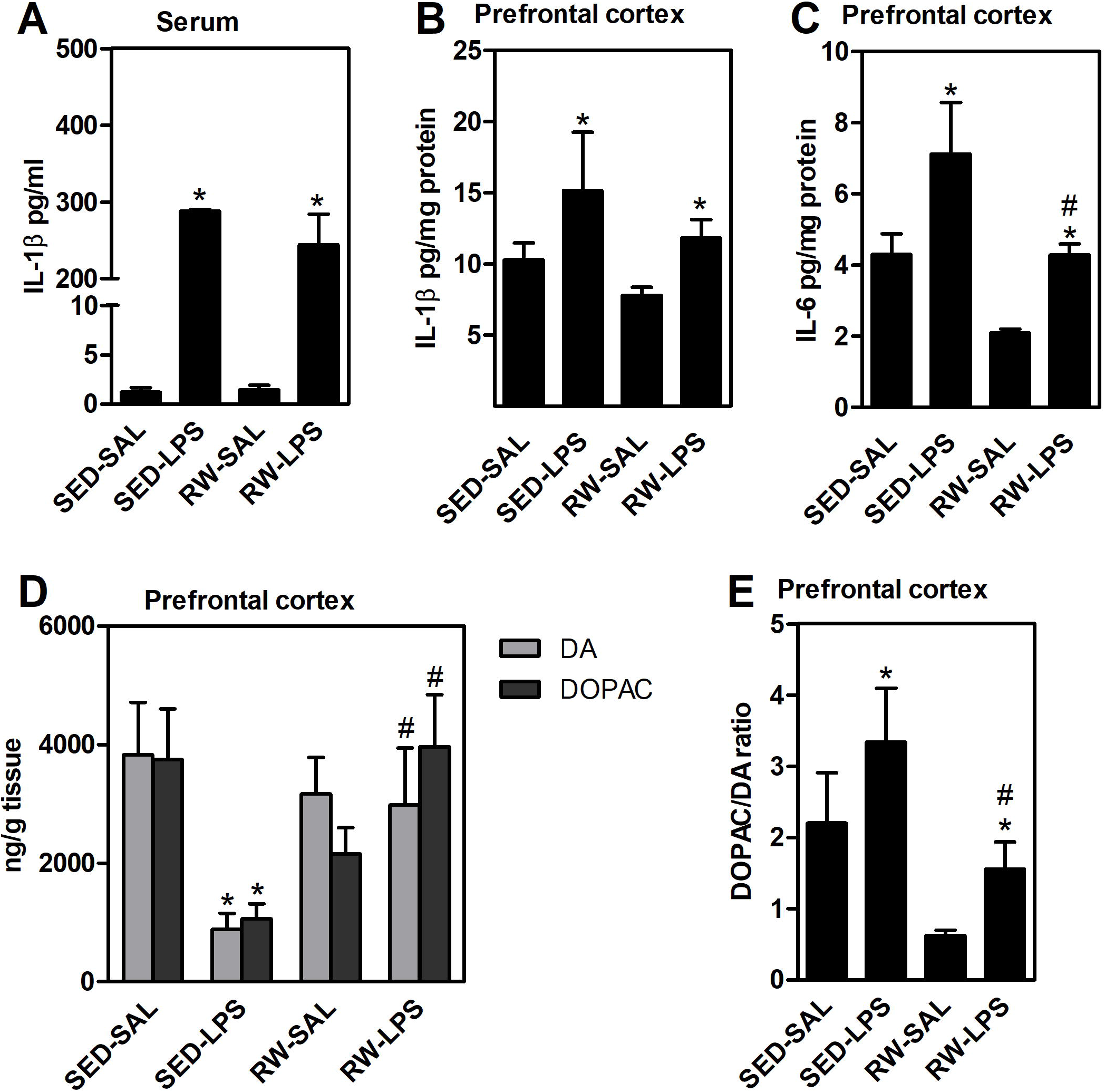
LPS (0.33 mg/kg, i.p.) induced systemic inflammation (A) and neuroinflammation (B-C). LPS reduced DA and DOPAC (D) and increased DOPAC/DA turnover (E) in the animals’ prefrontal cortex. The exercise partially reversed the increase in IL-6 (C) and DOPAC/DA turnover (E) in the prefrontal cortex. The recovery of DA and DOPAC levels was total (D). Data expressed as mean ± SEM for 9-10 independent experiments. * P<0.05 from SAL (ANOVA two-way). # P<0.05 from SED (ANOVA two-way). DA – dopamine. DOPAC – 3,4-dihydroxyphenylacetic acid. LPS – Lipopolysaccharide. RW – running wheel. SAL – saline. SED – sedentary.

LPS decreased DA [*F* (1,24) = 5.4, *p* < 0.05, η^2^ = 0.27, Figure 3D]; and DOPAC levels [*F* (1,27) = 11.2, *p* < 0.05, η^2^ = 0.29, Figure 3D] in the prefrontal cortex of animals, effects mitigated by exercise. Thus, LPS increased cortical dopaminergic turnover, decreased by exercise [*F* (1,25) = 5.9, *p*<0.05, η^2^ = 0.32, Figure 3E].

## DISCUSSION

This study demonstrated that LPS induced neuroinflammation and disruption of DA neurotransmission in the prefrontal cortex, as indicated by increased IL-1β and decreased DA levels, respectively, neuroinflammation-induced sickness behavior, and decreasing locomotor activity, food intake, self-care, and social interaction. Physical exercise decreased cortical DA turnover, maintaining DA levels in the prefrontal cortex, and attenuated neuroinflammation and sickness behavior in mice.

Decreased exploration of new environments (*e.g.*, open field) is a sensitive measure of sickness behavior [6]. Martin et al. [33, 48] did not observe the effects of exercise on LPS-induced mobility impairments in his experimental design: a rectangular arena of 0.57 m^2^. Here, exercise benefits to motor behavior were observed in a larger, circular arena, with 113 m^2^. The animal strains were different, C57BL6 for Martins et al. [33, 48], and Swiss in this study. These strains have different sensitivity to LPS, being higher in C57BL6 [49]. In common, a decrease in food intake and RW fatigue were observed in this work, and Martins et al. studies [33, 48], with no benefits from the previous exercise.

LPS also induces social withdrawal [42, 50]. We observed exercise-induced protection to self-care, anhedonia, social withdrawal in LPS-treated mice, and emotional components of behavior. Although there is no evidence of these latter effects, the literature is robust on the anxiolytic and antidepressant effects of exercise [51–53] in various animal models of depression and other neurological diseases [54, 55]. Sickness behavior is an adaptive reorganization of the host’s priorities during an infectious period, necessary for survival [33]. The benefits of exercise in attenuating, rather than disappearing, corroborate sickness behavior as a phenotype necessary for recovery.

The immune system is responsive to exercise in an intensity-dependent manner [56, 57]. High-intensity physical training increases the risk of upper respiratory tract infections, while mild-moderate intensity training protects against these infections [56, 58]. In this study, LPS increased serum levels of IL-1β and cortical levels of IL-1β and IL-6. Harden et al. (2008) [35] reported that both interleukin IL-1β and IL-6 act synergistically in the brain to induce sickness behavior. Godbout et al. (2005) also demonstrated that LPS increases IL-6 levels in the mouse brain [43]. However, the anti-inflammatory effects of exercise for LPS on the CNS are controversial. Martins et al. [33, 48] observed no effects of exercise (RW, 4-10 weeks) on increasing brain levels of IL-1β and IL-6 in LPS-treated C57BL6 mice. We used Swiss mice; light exercise partially reversed the cortical increase in IL-6 in LPS-treated animals. The greater sensitivity of the C57BL6 strain to LPS may explain this difference [49], making it impossible to detect differences. Evidence is robust in demonstrating anti-inflammatory effects of exercise on the brain of aged rodents [28] or models of Alzheimer’s [59] and Parkinson’s disease [60]. One mechanism is the negative regulation of TLR signaling [61]. TLR4 is the receptor for LPS that signals the cascade of proinflammatory cytokines. In humans, exercise decreases the expression of TLR4 in circulating monocytes [62, 63]. However, we did not see changes in serum and cortical levels of IL-1β.

Exercise protected from LPS-induced anhedonia, which is associated with low DA metabolism in the prefrontal cortex [64–66]. Here, LPS decreased DA concentration and its metabolite DOPAC in animals’ prefrontal cortex and induced anhedonia. Moreover, exercise reversed LPS-induced DA losses and DOPAC/DA ratio in the prefrontal cortex. DOPAC/DA ratio is an index of DA turnover and reflects synaptic DA terminals [29]. The exercise-induced decreased DOPAC/DA ratio suggests a decreased extracellular DA degradation at dopaminergic synapses, resulting in a higher probability of successful neurotransmission. Previous studies have also demonstrated protection from exercise against LPS-induced DA loss [60] and decreased cortical DOPAC/DA ratio in exercised animals [28, 52], associated with antidepressant and cognitive effects.

## CONCLUSION

Our study demonstrated that six weeks of light-intensity exercise attenuates sickness behavior and protects from dopaminergic impairment induced by neuroinflammation in mice. Previous studies also observed the effects of inflammatory cytokines on DA function, but the precise mechanism is still unknown. Our findings support physical exercise as a complementary strategy to recovery from behavioral impairments and exacerbated inflammation underlying neuroinflammatory conditions.

## DECLARATIONS

## ACKNOWLEDGEMENTS

This work was supported by the Coordenação de Aperfeiçoamento de Pessoal de Nível Superior–Brasil (CAPES)– Finance Code 1, Conselho Nacional de Desenvolvimento Científico e Tecnológico (CNPq) and Fundação de Amparo à Pesquisa e Inovação do Estado de Santa Catarina (FAPESC). We thank Dr. Rui Prediger (LEXDON/CCB/UFSC) and Dr. Alexandra Latini (LABOX/CCB/UFSC) for the use of scientific facilities. ASAJr, AL, and RP are CNPq fellow. ACBA is supported by CAPES/DS scholarship.

## ETHICAL PUBLICATION STATEMENT

We confirm that we have read the Journal’s position involving ethical publication and confirm that this report is accurate to the guidelines.

## AUTHOR CONTRIBUTIONS

**Study conception and design:** Alves, A.C.B.; Martins, R.P., Prediger, R.D.; Latini, A.; Aguiar Jr, AS.

**Acquisition of data:** Alves, A.C.B.; Pires, A.C.S.; Speck, A.E.; Scheffer, D.L.

**Analysis and interpretation of data:** Alves, A.C.B.; Pires, A.C.S.; Melo H.M.; Lopes, S.C.; Martins, R.P.

**Drafting of the manuscript:** Alves, A.C.B.; Aguiar Jr, AS.

**Critical revision:** Aguiar Jr, A.S.; Martins, R.P.; Alves, A.C.B. Latini, A.

## AVAILABILITY OF DATA AND MATERIALS

Any data used in this report that is not found in the reading can be available when requested to the corresponding author Aguiar Jr, A.S.

## CONSENT TO PARTICIPATE

All authors agreed to participate in this study and approved the final version.

## CONSENT FOR PUBLICATION

All authors have read and agreed to publish this manuscript.

## REFERENCES

1. Haroon E, Raison CL, Miller AH (2012) Psychoneuroimmunology meets neuropsychopharmacology: Translational implications of the impact of inflammation on behavior. Neuropsychopharmacology 37:137–162. https://doi.org/10.1038/npp.2011.205

2. Miller AH, Maletic V, Raison CL (2009) Inflammation and Its Discontents: The Role of Cytokines in the Pathophysiology of Major Depression. Biol Psychiatry 65:732–741. https://doi.org/10.1016/j.biopsych.2008.11.029

3. Maier SF, Watkins LR (1998) Cytokines for psychologists: Implications of bidirectional immune-to-brain communication for understanding behavior, mood, and cognition. Psychol Rev 105:83–107. https://doi.org/10.1037//0033-295x.105.1.83

4. Kiecolt-Glaser JK, McGuire L, Robles TF, Glaser R (2002) Emotions, Morbidity, and Mortality: New Perspectives from Psychoneuroimmunology. Annu Rev Psychol 53:83–107. https://doi.org/10.1146/annurev.psych.53.100901.135217

5. Shattuck EC, Muehlenbein MP (2015) Human sickness behavior: ultimate and proximate explanations. Am J Phys Anthropol 157:1–18. https://doi.org/10.1002/ajpa.22698

6. Konsman JP, Parnet P, Dantzer R (2002) Cytokine-induced sickness behaviour: Mechanisms and implications. Trends Neurosci 25:154–159. https://doi.org/10.1016/S0166-2236(00)02088-9

7. Berg BM, Godbout JP, Kelley KW, Johnson RW (2004) α-Tocopherol attenuates lipopolysaccharide-induced sickness behavior in mice. Brain Behav Immun 18:149–157. https://doi.org/10.1016/S0889-1591(03)00113-2

8. Larson SJ, Dunn AJ (2001) Behavioral effects of cytokines. Brain Behav Immun 15:371–387. https://doi.org/10.1006/brbi.2001.0643

9. Miller AH, Capuron L, Raison CL (2005) Immunologic influences on emotion regulation. Clin Neurosci Res 4:325–333. https://doi.org/10.1016/j.cnr.2005.03.010

10. Aubert A (1999) Sickness and behaviour in animals: a motivational perspective. Neurosci Biobehav Rev 23:1029–1036. https://doi.org/10.1016/S0149-7634(99)00034-2

11. Maes M (1995) Evidence for an immune response in major depression: A review and hypothesis. Prog Neuropsychopharmacol Biol Psychiatry 19:11–38. https://doi.org/10.1016/0278-5846(94)00101-M

12. Moylan S, Maes M, Wray NR, Berk M (2013) The neuroprogressive nature of major depressive disorder: Pathways to disease evolution and resistance, and therapeutic implications. Mol Psychiatry 18:595–606. https://doi.org/10.1038/mp.2012.33

13. Miller AH, Haroon E, Raison CL, Felger JC (2013) Cytokine Targets in the Brain: Impact on Neurotransmitters and Neurocircuits. Depress Anxiety 30:297–306. https://doi.org/10.1038/jid.2014.371

14. Mota CMD, Rodrigues-Santos C, Fernández RAR, et al. (2017) Central serotonin attenuates LPS-induced systemic inflammation. Brain Behav Immun 66:372–381. https://doi.org/10.1016/j.bbi.2017.07.010

15. Purnomo KI, Doewes M, Giri MKW, et al. (2017) Exercise Prevents Mental Illness. J Phys Conf Ser 180:1–6. https://doi.org/10.1088/1742-6596/755/1/011001

16. Barua CC, Haloi P, Saikia B, et al. (2018) Zanthoxylum alatum abrogates lipopolysaccharide-induced depression-like behaviours in mice by modulating neuroinflammation and monoamine neurotransmitters in the hippocampus. Pharm Biol 56:245–252. https://doi.org/10.1080/13880209.2017.1391298

17. Anderson G, Reiter RJ (2020) Melatonin: Roles in influenza, Covid-19, and other viral infections. Rev. Med. Virol.

18. Blatteis CM, Sehic E (1998) Cytokines and Fever. Ann N Y Acad Sci 840:608–618. https://doi.org/10.1111/j.1749-6632.1998.tb09600.x

19. Singhal G, Jaehne EJ, Corrigan F, et al. (2014) Inflammasomes in neuroinflammation and changes in brain function: A focused review. Front Neurosci 8:1–22. https://doi.org/10.3389/fnins.2014.00315

20. González H, Elgueta D, Montoya A, Pacheco R (2014) Neuroimmune regulation of microglial activity involved in neuroinflammation and neurodegenerative diseases. J Neuroimmunol 274:1–13. https://doi.org/10.1016/j.jneuroim.2014.07.012

21. Felger JC, Miller AH (2012) the Subcortical Source of Inflammatory Malaise. Front Neuroendocr 33:315–327. https://doi.org/10.1016/j.yfrne.2012.09.003.Cytokine

22. Chieffi S, Messina G, Villano I, et al. (2017) Neuroprotective effects of physical activity: Evidence from human and animal studies. Front Neurol 8:1–7. https://doi.org/10.3389/fneur.2017.00188

23. Hamer M, Chida Y (2008) Physical activity and risk of neurodegenerative disease: A systematic review of prospective evidence. Psychol Med 39:3–11. https://doi.org/10.1017/S0033291708003681

24. Middleton LE, Barnes DE, Lui L-Y, Yaffe K (2010) Physical Activity Over the Life Course and its Association with Cognitive Performance and Impairment in Old Age. J Am Geriatr Soc 58:1322–1326. https://doi.org/10.1038/jid.2014.371

25. Sofi F, Valecchi D, Bacci D, et al. (2011) Physical activity and risk of cognitive decline: A meta-analysis of prospective studies. J Intern Med 269:107–117. https://doi.org/10.1111/j.1365-2796.2010.02281.x

26. Yaffe K, Fiocco AJ, Lindquist K, et al. (2009) Predictors of maintaining cognitive function in older adults. Neurology 72:2029–2035. https://doi.org/10.1212/WNL.0b013e3181a92c36

27. Littlefield AM, Setti SE, Priester C, Kohman RA (2015) Voluntary exercise attenuates LPS-induced reductions in neurogenesis and increases microglia expression of a proneurogenic phenotype in aged mice. J Neuroinflammation 12:1–12. https://doi.org/10.1186/s12974-015-0362-0

28. Dallagnol KMC, Remor AP, da Silva RA, et al. (2017) Running for REST: Physical activity attenuates neuroinflammation in the hippocampus of aged mice. Brain Behav Immun 61:31–35. https://doi.org/10.1016/j.bbi.2016.07.159

29. Aguiar AS, Lopes SC, Tristão FSM, et al. (2016) Exercise Improves Cognitive Impairment and Dopamine Metabolism in MPTP-Treated Mice. Neurotox Res 29:118–125. https://doi.org/10.1007/s12640-015-9566-4

30. Lee H, Nagata K, Nakajima S, et al. (2018) Intermittent intense exercise protects against cognitive decline in a similar manner to moderate exercise in chronically stressed mice. Behav Brain Res 345:59–64. https://doi.org/10.1016/j.bbr.2018.01.017

31. Li B, Liang F, Ding X, et al. (2019) Interval and continuous exercise overcome memory deficits related to β-Amyloid accumulation through modulating mitochondrial dynamics. Behav Brain Res 376:1–10. https://doi.org/10.1016/j.bbr.2019.112171

32. Davis JM, Murphy EA, Brown AS, et al. (2004) Effects of moderate exercise and oat β-glucan on innate immune function and susceptibility to respiratory infection. Am J Physiol - Regul Integr Comp Physiol 286:366–372. https://doi.org/10.1152/ajpregu.00304.2003

33. Martin SA, Pence BD, Greene RM, et al. (2013) Effects of voluntary wheel running on LPS-induced sickness behavior in aged mice. Brain Behav Immun 29:113–123. https://doi.org/10.1016/j.bbi.2012.12.014

34. Harden LM, du Plessis I, Poole S, Laburn HP (2006) Interleukin-6 and leptin mediate lipopolysaccharide-induced fever and sickness behavior. Physiol Behav 89:146–155. https://doi.org/10.1016/j.physbeh.2006.05.016

35. Harden LM, Plessis I du, Poole S, Laburn HP (2008) Interleukin (IL)-6 and IL-1β act synergistically within the brain to induce sickness behavior and fever in rats. Brain Behav Immun 22:838–849. https://doi.org/10.1016/j.bbi.2007.12.006

36. Criswell DS, Henry KM, DiMarco NM, Grossie VB (2004) Chronic exercise and the proinflammatory response to endotoxin in the serum and heart. Immunol Lett 95:213–220. https://doi.org/10.1016/j.imlet.2004.07.012

37. Chen HI, Hsieh SY, Yang FL, et al. (2007) Exercise training attenuates septic responses in conscious rats. Med Sci Sports Exerc 39:435–442. https://doi.org/10.1249/mss.0b013e31802d11c8

38. Koo JH, Kwon IS, Kang EB, et al. (2013) Neuroprotective effects of treadmill exercise on BDNF and PI3-K/Akt signaling pathway in the cortex of transgenic mice model of Alzheimer’s disease. J Exerc Nutr Biochem 17:151–160. https://doi.org/10.5717/jenb.2013.17.4.151

39. Aguiar Jr. AS, Tristão FSM, Amar M, et al. (2014) Six weeks of voluntary exercise don’t protect C57BL/6 mice against neurotoxicity of MPTP and MPP^+^. Neurotox Res 25:. https://doi.org/10.1007/s12640-013-9412-5

40. Cardoso L, Nery T, Goncalves M, et al. (2020) Caffeine decreases neuromuscular fatigue in the lumbar muscles - a randomized blind study. medRxiv 2020.06.08.20122531. https://doi.org/10.1101/2020.06.08.20122531

41. Henry CJ, Huang Y, Wynne AM, Godbout JP (2009) Peripheral Lipopolysaccharide (LPS) challenge promotes microglial hyperactivity in aged mice that is associated with exaggerated induction of both proinflammatory IL-1β and anti-inflammatory IL-10 cytokines.pdf. Brain Behav Immun 23:309–317. https://doi.org/10.1016/j.bbi.2008.09.002.Peripheral

42. Henry CJ, Huang Y, Wynne A, et al (2008) Minocycline attenuates lipopolysaccharide (LPS)-induced neuroinflammation, sickness behavior, and anhedonia. J Neuroinflammation 5:1–14. https://doi.org/10.1186/1742-2094-5-15

43. Godbout JP, Chen J, Abraham J, et al. (2005) Exaggerated neuroinflammation and sickness behavior in aged mice following activation of the peripheral innate immune system. FASEB J 19:1329–1331. https://doi.org/10.1096/fj.05-3776fje

44. Seibenhener ML, Wooten MC (2015) Use of the open field maze to measure locomotor and anxiety-like behavior in mice. J Vis Exp 1–6. https://doi.org/10.3791/52434

45. Isingrini E, Camus V, Le Guisquet AM, et al. (2010) Association between repeated unpredictable chronic mild stress (UCMS) procedures with a high fat diet: A model of fluoxetine resistance in mice. PLoS One 5:. https://doi.org/10.1371/journal.pone.0010404

46. Kaidanovich-Beilin O, Lipina T, Vukobradovic I, et al. (2010) Assessment of social interaction behaviors. J Vis Exp 0:1–6. https://doi.org/10.3791/2473

47. Lowry (1951) Lowry Protein Assay. J Biol Chem

48. Martin SA, Dantzer R, Kelley KW, Woods JA (2014) Voluntary Wheel Running Does not Affect Lipopolysaccharide-Induced Depressive-Like Behavior in Young Adult and Aged Mice. Neuroimmunomodulation 21:52–63. https://doi.org/10.1038/jid.2014.371

49. Dias RS, Carmo LS, Heneine LGD, et al (2009) The use of mice as animal model for testing acute toxicity (LD-50) of toxic shock syndrome toxin. Arq Bras Med Veterinária e Zootec. https://doi.org/10.1590/s0102-09352009000100024

50. Kirsten TB, Taricano M, Maiorka PC, et al. (2010) Prenatal lipopolysaccharide reduces social behavior in male offspring. Neuroimmunomodulation 17:240–251. https://doi.org/10.1159/000290040

51. Aguiar AS, Latini A (2018) Treating depression with exercise: An immune perspective

52. Cunha MP, Oliveira Á, Pazini FL, et al. (2013) The antidepressant-like effect of physical activity on a voluntary running wheel. Med Sci Sports Exerc 45:851–859. https://doi.org/10.1249/MSS.0b013e31827b23e6

53. Aguiar AS, Stragier E, da Luz Scheffer D, et al. (2014) Effects of exercise on mitochondrial function, neuroplasticity and anxio-depressive behavior of mice. Neuroscience 271:56–63. https://doi.org/10.1016/j.neuroscience.2014.04.027

54. Russo-Neustadt A, Ha T, Ramirez R, Kesslak JP (2001) Physical activity-antidepressant treatment combination: Impact on brain-derived neurotrophic factor and behavior in an animal model. Behav Brain Res. https://doi.org/10.1016/S0166-4328(00)00364-8

55. Brené S, Bjørnebekk A, Åberg E, et al (2007) Running is rewarding and antidepressive. Physiol Behav. https://doi.org/10.1016/j.physbeh.2007.05.015

56. Nieman DC (1994) Exercise, infection, and immunity. In: International Journal of Sports Medicine

57. Laddu DR, Lavie CJ, Phillips SA, Arena R (2020) Physical activity for immunity protection: Inoculating populations with healthy living medicine in preparation for the next pandemic. Prog. Cardiovasc. Dis.

58. Scheffer D da L, Latini A (2020) Exercise-induced immune system response: Anti-inflammatory status on peripheral and central organs. Biochim. Biophys. Acta - Mol. Basis Dis.

59. Parachikova A, Nichol KE, Cotman CW (2008) Short-term exercise in aged Tg2576 mice alters neuroinflammation and improves cognition. Neurobiol Dis. https://doi.org/10.1016/j.nbd.2007.12.008

60. Wu SY, Wang TF, Yu L, et al. (2011) Running exercise protects the substantia nigra dopaminergic neurons against inflammation-induced degeneration via the activation of BDNF signaling pathway. Brain Behav Immun 25:135–146. https://doi.org/10.1016/j.bbi.2010.09.006

61. Abbasi A, Hauth M, Walter M, et al. (2014) Exhaustive exercise modifies different gene expression profiles and pathways in LPS-stimulated and un-stimulated whole blood cultures. Brain Behav Immun. https://doi.org/10.1016/j.bbi.2013.10.023

62. Lancaster GI, Khan Q, Drysdale P, et al. (2005) The physiological regulation of toll-like receptor expression and function in humans. J Physiol. https://doi.org/10.1113/jphysiol.2004.081224

63. Flynn MG, McFarlin BK, Phillips MD, et al. (2003) Toll-like receptor 4 and CD14 mRNA expression are lower in resistive exercise-trained elderly women. J Appl Physiol. https://doi.org/10.1152/japplphysiol.00359.2003

64. Espejo EF (2003) Prefrontocortical dopamine loss in rats delays long-term extinction of contextual conditioned fear, and reduces social interaction with out affecting short-term social interaction memory. Neuropsychopharmacology 28:490–498. https://doi.org/10.1038/sj.npp.1300066

65. Wilson CA, Koenig JI (2014) Social interaction and social withdrawal in rodents as readouts for investigating the negative symptoms of schizophrenia. Eur Neuropsychopharmacol 24:759–773. https://doi.org/10.1016/j.euroneuro.2013.11.008.Social

66. Zhang J, He ZX, Wang LM, et al. (2019) Voluntary wheel running reverses deficits in social behavior induced by chronic social defeat stress in mice: Involvement of the dopamine system. Front Neurosci 13:1–11. https://doi.org/10.3389/fnins.2019.00256

